# A Spectral De-mixing Model for Triplex *In Vivo* Imaging of Optical Coherence Tomography Contrast Agents

**DOI:** 10.1101/846170

**Authors:** Edwin Yuan, Peng Si, Saba Shevidi, Adam de la Zerda

## Abstract

The ability to detect multiple contrast agents simultaneously would greatly enhance Optical Coherence Tomography (OCT) images, providing nuanced biological context to physiological structures. However, previous OCT contrast agent work has been limited to scenarios where only a single contrast agent could be robustly detected within each voxel. We present a novel spectroscopic technique for de-mixing the spectral signal from multiple OCT contrast agents within a single voxel. We validate our technique *in vitro* and also demonstrate *in vivo* imaging of three spectrally distinct gold nanobipyramids, trafficking within the lymphatic system of a live mouse. This approach opens the door to a much broader range of pre-clinical and clinical OCT applications where multiplexed labeling is desirable.

## Main Results

Optical Coherence Tomography (OCT) is an interferometric imaging technique that allows for non-invasive 3D tomography of the scattering properties of tissue at micron resolution, with improved depth sectioning and imaging depth compared to incoherent imaging^1^. A significant limitation of OCT, however, is the lack of multiplexed biological labeling techniques that are available to other imaging techniques (e.g. fluorescent markers), which enable localization of cell types, patterns of protein expression, and imaging of lymph/blood vessels^2^. The majority of previous work has focused on use of a single contrast agent, such as absorptive dyes^3^, microbeads^4,5,6^, and gold nanoparticles^7,8,9^. Two-plex imaging of gold nanoparticles has been demonstrated, but only under the assumption that each voxel contained a single type of contrast agent^10,11^. The biological applications of contrast-enhanced OCT techniques would be greatly expanded by the ability to distinguish multiple species of the contrast agent^12^ within a single voxel.

In this work, we demonstrate a novel spectroscopic processing technique that enables highly multiplexed OCT contrast agent imaging, utilizing spectrally distinct gold nanoparticles. The technique poses the Short Time Fourier Transform (STFT) spectra measured within each voxel as a sum of spectra measured from individual contrast agents, solving for their proportions through numerical optimization. To the best of our knowledge, the de-mixing framework established in this work is the first to enable “true” de-mixing, in which the contrast agent proportions within each voxel can be predicted. As a proof-of-concept, we demonstrate the first instance of simultaneously imaging three OCT contrast agents.

The contrast agents used in this work are gold nanobipyramids (GNBPs)^13^, gold nanoparticles resembling two elongated pyramids fused at their bases. GNBPs have highly tunable aspect ratio, strong scattering cross-sections^11^, and surface plasmon resonances in the second near infrared window (1100~1700 nm), making them ideal OCT contrast agents for *in vivo* imaging. We synthesized 3 different GNBPs with dimensions of 137 nm × 22 nm, 154 × 23 nm and 183 nm × 25 nm (Figure 1a), with center wavelengths at 1225, 1315, and 1450 nanometers, and full width half maximum of 45 nm, 50 nm, 80 nm (Figure 1b), henceforth named GNBP_1225_, GNBP_1315_, and GNBP_1450_.

**Figure 1.**
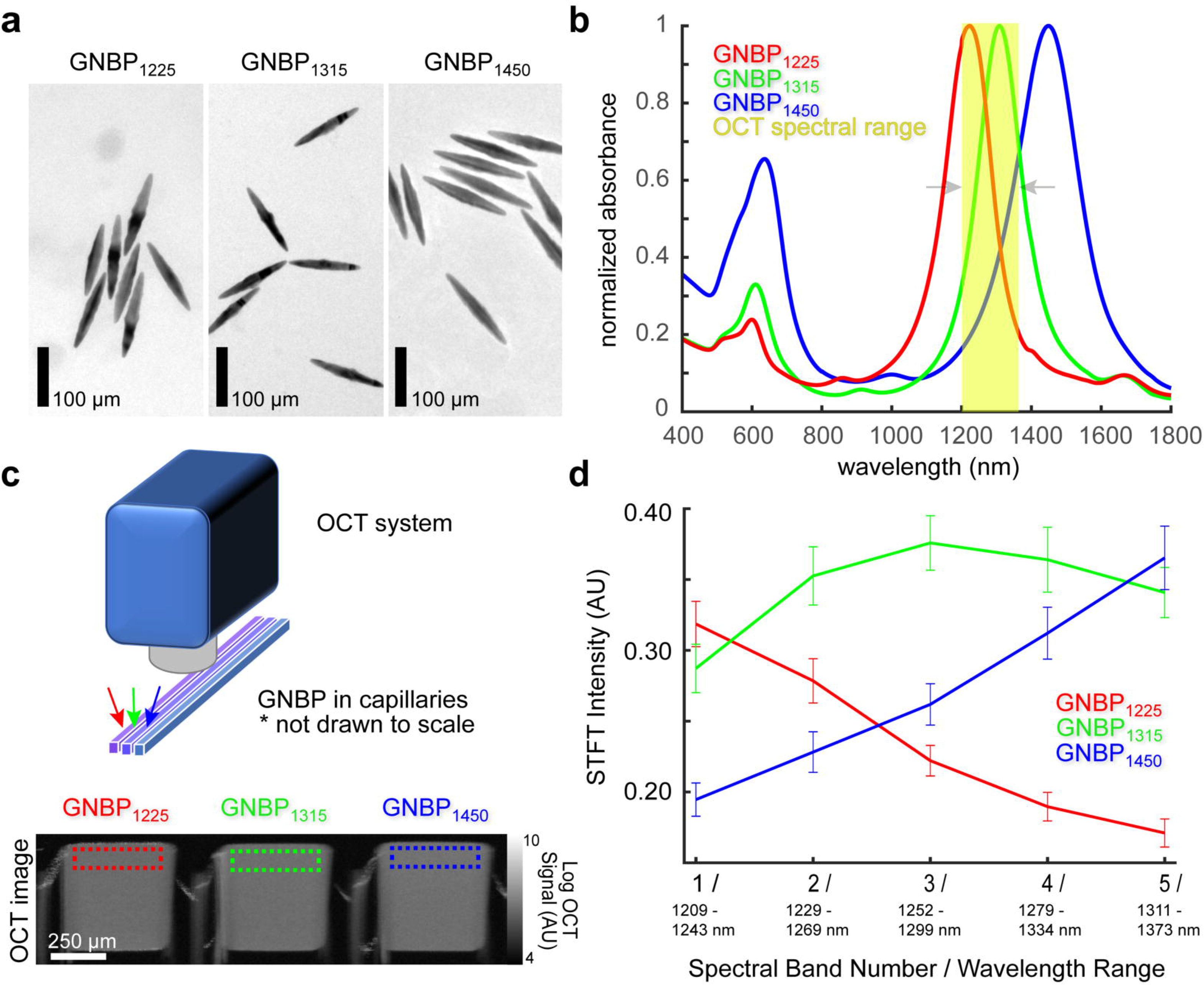
OCT Imaging of GNBPs. **a** Three species of GNBPs with different peak wavelengths, 1225 nm, 1315 nm, and 1450 nm, imaged in a transmission electron microscope. Scale bar: 100 nm. **b** UV-Vis-NIR measured absorbance of GNBP_1225_, GNBP_1315_, and GNBP_1450_. The spectrometer range of the OCT system (Thorlabs Telesto-II) utilized in this work is shown in yellow for comparison. **c** Top: Camera image of GNBP loaded into glass capillaries and imaged in OCT system. Bottom: OCT image of 3 glass capillaries containing GNBP_1225_, GNBP_1315_, and GNBP_1450_. The dotted boxes show the ROIs for quantitative spectral analysis. **d** Average spectra of GNBPs in the dotted box ROIs from the OCT image in c. Spectra are computed as a five-band STFT of the OCT interferogram. Error bars show the standard deviation of each spectral band.

To demonstrate the signal de-mixing capability of our spectroscopic processing techniques *in vitro*, we imaged GNBP_1225_, GNBP_1315_, and GNBP_1450_ in glass capillaries (Figure 1c, S1a) alongside 50/50 mixtures of every pair of particle types. We employ the discrete short time fourier transform (STFT), dividing the spectral bandwidth of the OCT system into 5 bands of equal size of about Δλ = 44 nm, with overlap of 26.5 nm, reducing the axial resolution from 3.5 um to 11.9 um in water^14^. The STFT yields an image in which each pixel contains a 5-dimensional vector, corresponding to the spectrum of the light scattered from that position. The STFT spectrum of each pure particle species resembles the UV-Vis absorbance measurements (Figure 1b,1c), with low variance (less than 15%) in each spectral band after B-scan averaging.

We visualize the separability of the STFT spectra of different GNBP species using principle component analysis (PCA)^15^ for dimensionality reduction. We apply PCA to the 5-dimensional STFT spectra of each pixel, mapping it to a lower, 3-dimensional space, which can be represented as an RGB image (Figure S1a). Particularly, when we plot the 2 leading PCA components on a scatter plot (Figure S1b), the spectra of each of the 3 pure GNBP species lies within an easily distinguishable cluster, with the secondary clusters representing 50/50 mixed particles lying in between.

We formulate a de-mixing model for the STFT spectra of each pixel as a decomposition of the 3 reference GNBP species, allowing us to infer the relative concentration and, thus, proportion of each GNBP species. Our model utilizes the non-negative least squares algorithm to arrive at solutions which optimally satisfy the de-mixing model. We evaluate this predictive performance by measuring 10 capillaries containing the pure particle species (3 leftmost columns in Figure 2a), as well as various known mixtures of 2 and 3 GNBP species. The OCT image of the capillaries gives no indication that each capillary contains a spectrally unique combination of GNBP (Figure 2a). Our model, however, reveals the individual de-mixed spectra (Figure 2b), mapped to an RGB color scheme, as well as the merged channels (Figure 2c). We also plot both the predicted GNBP proportions and the ground truth GNBP proportions (Figure 2d). Our results demonstrate an error in the proportions of each GNBP species of no more than 11% (Figure 2e). The de-mixing error is notably higher for the pure GNBP_1315_ (11%) compared to the pure GNBP_1225_ (5%) and GNBP_1450_ (5%), likely due its spectral peak lying between those of the other two GNBPs.

**Figure 2.**
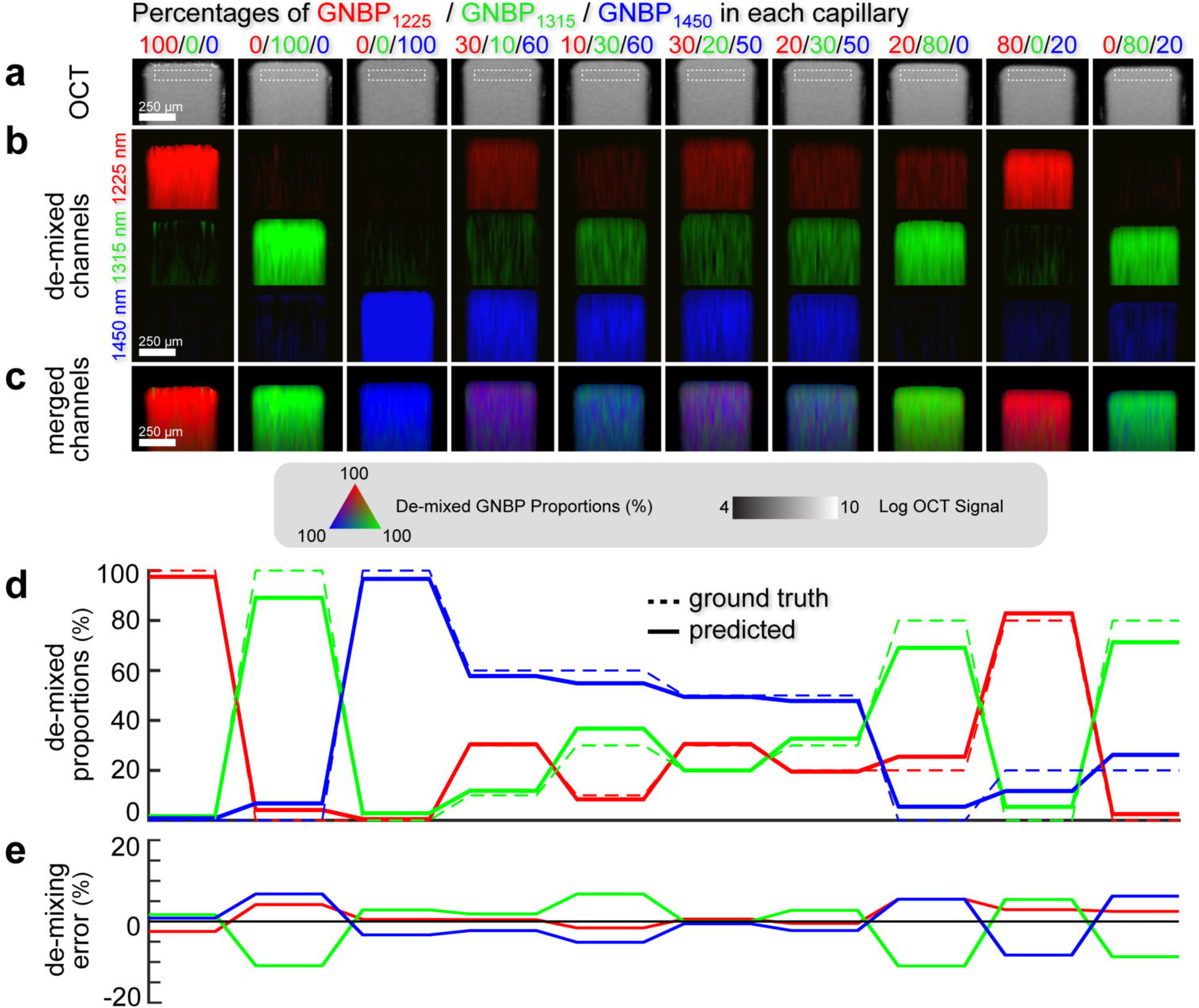
De-mixing of Spectral Signals from GNBPs. **a** OCT image of glass capillaries containing various mixtures of GNBPs with combined concentration 0.167 nM. From left to right: Capillaries 1-3 contain pure GNBP_1225_, pure GNBP_1315_, and pure GNBP_1450_. Capillaries 4-7 contain mixtures of 3 different GNBPs. Capillaries 8-10 contain mixtures of two GNBPs. Spectral signal is calculated from ROIs within dotted white boxes at the top of each capillary. Scale bar: 250 μm. **b** De-mixed GNBP channels from **a**. Red: 1225 nm GNBP channel. Blue: 1315 nm GNBP channel. Green: 1450 nm GNBP channel. Scale bar: 250 μm. **c** Merged RGB channels from **b**. Scale bar: 250 micrometers. **d** Proportions of each GNBP channel from **c**, for each of the 10 capillaries. The proportions for each capillary sum to 1. The solid line shows the predicted proportions from the de-mixing algorithm, while the dotted line shows the actual proportions. **e** De-mixing error, the difference between the solid and dotted lines in **d**, for each channel and each capillary.

Next, we utilized our ability to simultaneously image 3 nanoparticles to study the spatial dynamics of lymphatic drainage *in vivo*. In the ear of an anesthetized mouse, we made 3 injections of 0.1 μL each, of 5nM GNBP_1225_, GNBP_1315_, and GNBP_1450_, with imaging 15 minutes (Figure 3) and 60 minutes after injection (Figure S2). Using the spectral processing techniques described above, we produced the de-mixed OCT image of a 6×6×2.5 mm volume. We only show voxels with sufficient speckle variance, in order to isolate blood vessels, and lymphatic vessels with flowing GNBPs. The ratio of colors in each voxel indicates the proportion of each particle species, while the brightness of the colors indicates the OCT signal intensity. We observe 3 outlined injection sites where a high concentration of GNBP are diffusing in subcutaneous tissue. A large network of lymphatic vessels is observed in each proximity, trafficking GNBP to a downstream lymph node. We note several junctions where GNBPs of different species merge and two lymphatic vessels, labeled 1 and 2, where this is especially salient.

**Figure 3.**
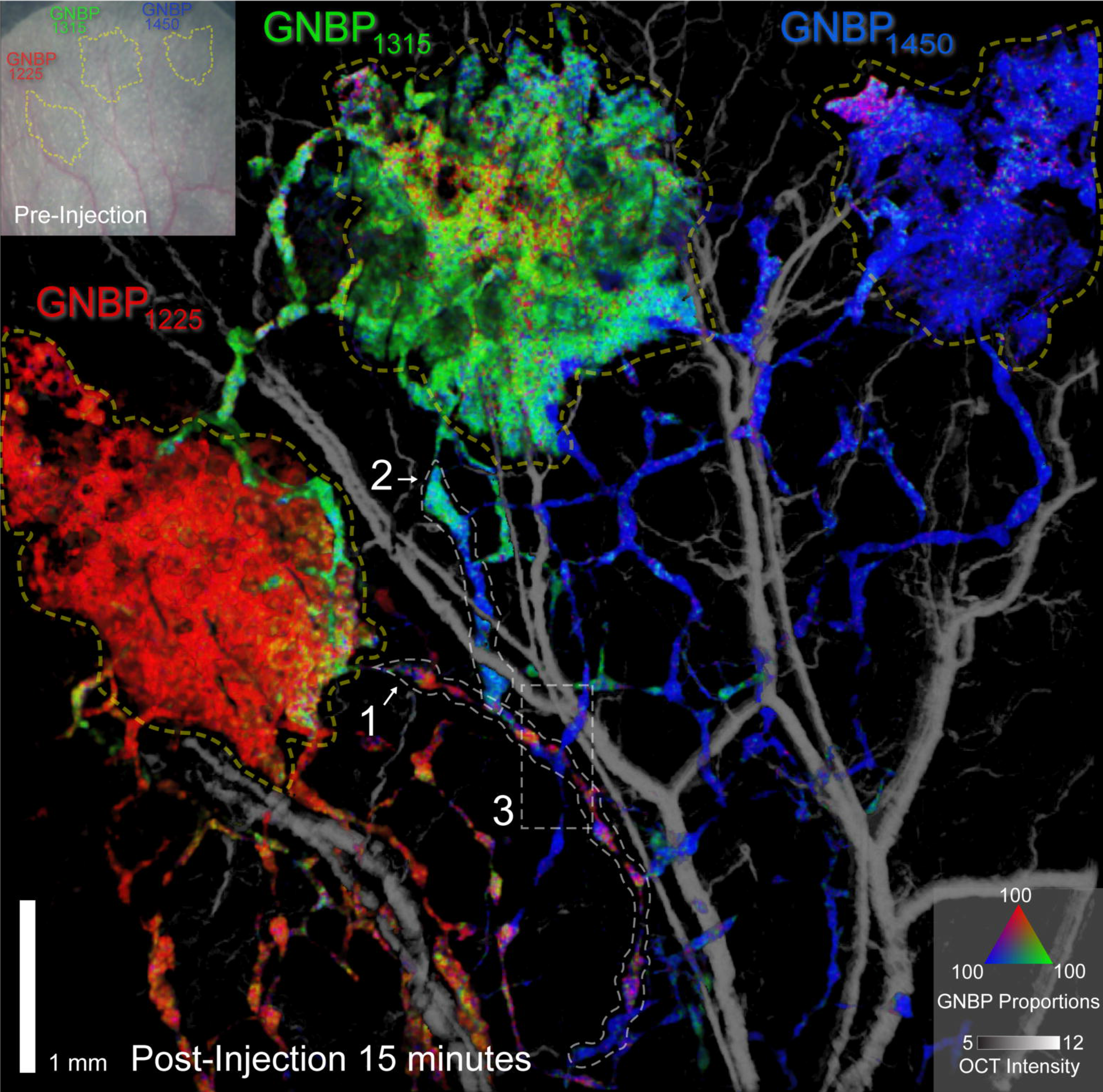
*In vivo* imaging of 3 GNBP species in mouse ear. Inset shows mouse ear pre-injection, with the 3 injection sites shown as dotted yellow outlines. The figure shows the speckle variance gated OCT image of the mouse ear 15 minutes postinjection. Lymphatic vessels are displayed in color, while blood vessels are shown in grayscale. For lymphatic vessels, the RGB color proportions indicate the proportions of GNBP_1225_, GNBP_1315_, and GNBP_1450_ predicted by the de-mixing algorithm. For both lymphatic and blood vessels, the intensity indicates the OCT signal intensity. The injection sites are outlined using a dotted yellow line. Lymphatic vessels 1 and 2, where significant mixing of nanoparticles take place, are outlined using white dotted lines. Scale bar: 1 mm.

Animal imaging leads to additional uncertainty in the spectrum arriving at the top of each lymphatic vessel, due to spectrally dependent absorption and scattering within tissue. A spectral calibration technique is developed to account for this uncertainty, by normalizing the spectra of certain voxels, with a high concentration of a single GNBP type, to the reference GNBP spectra. We address the accuracy of the spectrally calibrated *in vivo* de-mixing in Figure S3. The results (Table S1) show that the average de-mixing error for the pure particle species is 9.2% *in vivo*, compared to 7% for the *in vitro* experiments (Figure 2d).

We more closely analyze lymph 1 at 15 minutes post-injection (Figure 4a), by dividing it into 9 sections, and showing the proportion of GNBPs for each section along its length (Figure 4c). Lymph 1 emerges near the GNBP_1225_ nm injection site, and thus contains a majority of these nanoparticles for most of its length. At section 4, lymph 1 merges with an upstream vessel containing GNBP_1450_, leading to elevated proportions of GNBP_1450_ in sections 4,6, and 7. However, when observing lymph 1 at 60 minutes after injection (Figure 4b), we see a dramatic change in the color of the lymphatic vessel, indicating a change in the composition of GNBPs (Figure S4). It now contains a majority of GNBP_1315_.This can be explained by the fact that one branch of the lymph network from the GNBP_1315_ injection site terminates close to section 1 of lymph 1.

**Figure 4.**
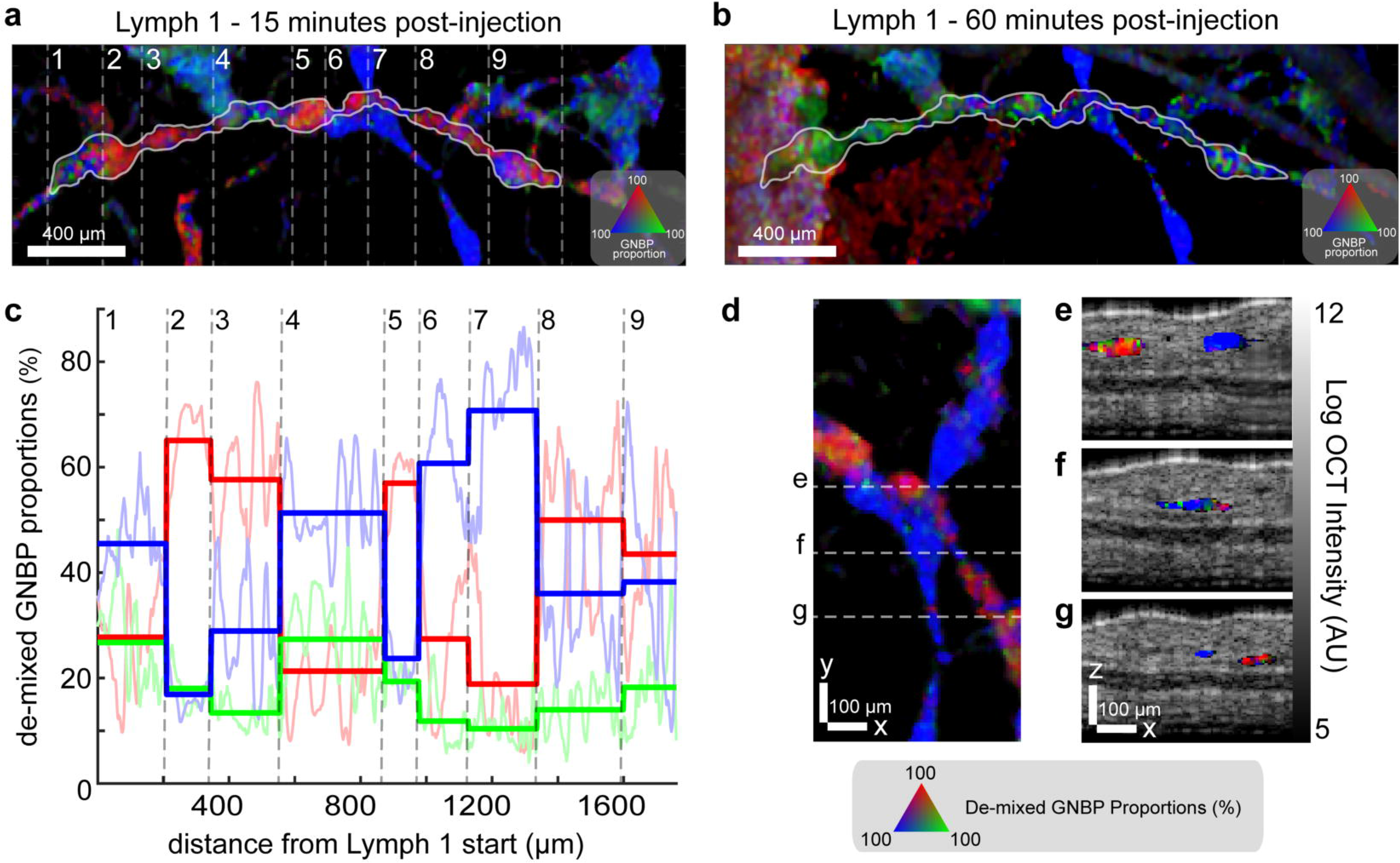
Analysis of the GNBP proportions in lymphatic vessels. **a** De-mixed image of lymph vessel 1 from **Figure 3**, 15 minutes post-injection, highlighted by a white boundary. The vessel is segmented into several sections by dotted lines. **b** De-mixed image of lymph 1 at 60 minutes post-injection, showing substantial change in the proportion of GNBPs compared to **a**. For **a,b** scale bar: 400 μm. **c** De-mixed proportions of GNBPs in lymph 1, taken as the mean of each section within the dotted boundaries. The saturated lines indicate averaged de-mixing proportions within each section, while dimmed lines show raw data prior to averaging. **d** De-mixed image of two lymph vessels, from ROI 3 in **Figure 3**, carrying predominantly different GNBPs. **e,f,g** Traverse OCT cross-sections of lymph vessels shown by dotted lines in **d**. Without knowledge of the de-mixed GNBPs, the vessels appear to intersect, but the de-mixing shows that the GNBP_1225_ and GNBP_1450_ flows in fact do not merge. Scale bars: 100 μm.

Lymph 2 (Figure S5) exhibits the presence of valves, which prevent lymphatic backflow^16^, as seen in the sharp boundary between section 1, containing primarily GNBP_1315_, and section 2, containing GNBP_1450_ (Figure S5a). Interestingly, in spite of its proximity to the GNBP_1315_ injection site, lymph 2 maintains an abundance of GNBP_1450_ even 60 minutes post-injection (Figure S5b). This suggests that pulsing lymphatic flow, often described in literature^17^, was insufficient to completely overturn the proportions of GNBP in the lymph vessel.

Finally, we examine a lymphatic junction in region 3 (Figure 3), which shows the apparent intersection of two lymphatic vessels, one carrying primarily GNBP_1225_ and other GNBP_1450_. The two vessels appear to directly contact each other (Figure 4e,f,g), but our multiplexing capability shows that they contain separate, non-intersecting channels of lymphatic flow. This illustrates the surprising fact that even lymphatic vessels in close proximity to each other can contain drainage from spatially distinct parts of tissue.

In summary, this work presents, for the first time, the ability to de-mix OCT contrast agent signals both *in vitro* and *in vivo* within a single voxel, and the first demonstration of triplex OCT imaging. Future work will focus on conjugation of GNBPs, incorporating speckle reduction^18^, and utilizing wider-bandwidth OCT light sources, which allow for greater multiplexing with less reduction in axial resolution^19^. Building on previous OCT work demonstrating tracking of cancer cells^20^, lymph biomarkers^21^, and conjugation of antibodies to gold nanorods^10^, there are numerous pre-clinical and clinical opportunities enabled by this work. An example would be imaging the spatiotemporal dynamics of various immunocytes such as CD8+ T cells, NK cells, and tumor-associated macrophages in the tumor microenvironment of a live animal model, and characterizing their cellular response to immunotherapies. Such a platform is posed to bring many of the most powerful investigative tools of multiplexed molecular labeling to OCT.

## Methods

### OCT setup

All results from this work were acquired using the Thorlabs Telesto-II OCT system (ThorLabs, Newton, NJ). The light source for the Telesto-II system consists of two superluminescent diodes, and the spectrometer has a spectral range from 1208.69 to 1372.50 nm, providing axial resolution of 3.96 μm in water. Additionally, the spectrometer has 1024 pixels, and was acquiring OCT a-lines at a rate of 91 kHz. All images were acquired with a LSM03 lens (ThorLabs, Newton, NJ), which provided a lateral resolution of 11.59 μm FWHM, and a depth of field (DOF) of 270 μm. Additionally, all images have an isotropic lateral pixel size of 6 μm and were taken with 50 B-scan averages.

### Creation of Gold Nanobipyramids

The gold nanobipyramids (GNBPs) were synthesized using a reported protocol^13^ with some modifications. In general, the aspect ratio of the particles can be controlled by altering the amount of surfactant and the amount of gold seed used in producing the particles. The gold seeds were prepared by mixing 4 mL of 0.5 mM HAuCl_4_, 4 mL of 95 mM cetyltrimethylammonium chloride (CTAC) and 72 μL of 250 mM HNO_3_, followed by rapidly injecting 100 μL of 50 mM ice-cold NaBH4 solution containing 50 mM NaOH under vigorous stirring. After the mixture was stirred for 1 min, 16 μL of 1 M citric acid was added to the solution and the cap of the reaction vial was tightly closed. The vial containing this seed solution was then placed in a water bath of 80 °C for aging. After 1 h, the seed solution became reddish, indicating an increase in seed size, and it was removed from the water bath. The growth solutions of GNBP_1225_, GNBP_1315_ and GNBP_1450_ were prepared by dissolving 48.5, 42.5 and 35 mg CTAB respectively in 400 mL of 140 mM CTAC solution, followed by adding 4 mL of 25 mM HAuCl_4_, 1.2 mL of 10 nM silver nitrate, and 4 mL of 0.4 M ethanol solution of 8-hydroxyquinoline. Then 2 mL of the aged seed solution was injected into the growth solution, which was gently stirred for 10 s and then placed in a water bath at 45 °C for 15 min. After that, 3 mL of 0.4 M ethanol solution of 8-hydroxyquinoline was added to the growth solution, which was kept in the 45 °C water bath for another 1.5 h. After the reaction, the growth solution turned dark blue. To collect the resulting GNBPs, the growth solution was transferred to 250 mL Nalgene centrifuge bottles, which were centrifuged at 30,000 g for 30 min. The supernatant was discarded, and the obtained pellet was washed by double distilled water twice. To PEGylate the nanoparticles, 0.5 nM GNBP was incubated with 0.5 mg/mL solution of mPEG-SH 5000 overnight at 4 °C.

The NIR spectra of the GNBPs were measured by UV-Vis-NIR spectrometer (Jasco Model V670). The surface morphology and dimensions of the GNBPs were characterized by transmission electron microscopy (FEI Tecnai G2 F20 X-TWIN).

### Spectral De-mixing Algorithm

The essential idea underlying the spectral de-mixing algorithm is that the square of the observed spectrum in each pixel is a sum of squares of the spectra of each of the 3 GNBP’s, weighted by their concentrations.

The coherent scattering field *E* from numerous particles in a voxel follows Rayleigh’s distribution^22^, which describes the average magnitude of the sum of *n* randomly oriented phasors (each representing a single particle) *E_p_*.

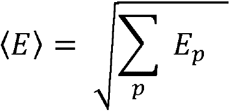

Experimentally, we verify the correctness of this description by measuring the scattering of GNBP in a capillary as a function of the concentration, which follows a square root law as expected (Figure S6). The short time fourier transform measures the scattering field in each voxel, as a function of wavenumber, which we now denote as 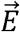. We correct the measured scattering field 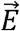 for the wavenumber dependent OCT sensitivity fall-off (Figure S7), although, in practice, this is a small effect that has a negligible outcome on the de-mixing results. Each GNBP_n_ also has a distinct scattering spectrum, which we denote as 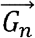, and a distinct concentration *c_n_*.

Finally, there are two additional factors which modify the scattering spectrum 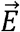. The first is chromatic aberration 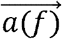, which depends on the position of the focus in the OCT image (Figure S8). The second is a depth-dependent modification to the electric field due to absorption and scattering above the current pixel position. The modified electric field reaching a particular pixel is thus 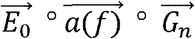.

We formulate the scattering spectrum from every pixel as:

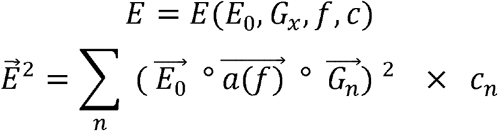

where ° refers to the element-wise dot product of two vectors. We want to decompose this measurement of 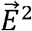 in terms of the reference GNBP_n_ measured in capillaries.

For the reference measurement *E^R^* performed in the capillary, we denote particular values *f^R^*, 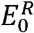, and 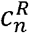.

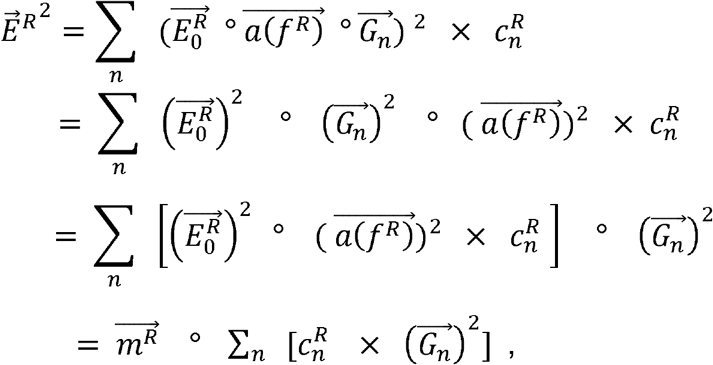

, where 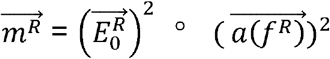 is a vector accounting for both the modification to the GNBP electric field and chromatic aberration.

For simplicity we set 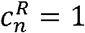 for the reference measurement, so:

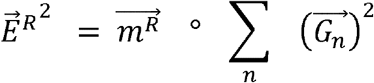

We can also write the measurement of each pixel in our *in vivo* volume as *E^p^*

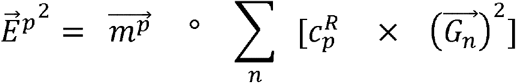

In order to equate *E^p^*, the pixel value in our acquired volume with the measured electric field of the reference measurement, *E^R^*, we must ensure that both measurements are subject to the same spectral modification due to absorption, scattering, and chromatic aberration. Thus, we transform the acquired volume via the factor 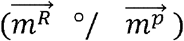, obtaining:

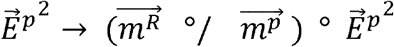

Equating the aberration corrected *E^p^* with the reference spectrum *E^R^*, with additional weights, *w_n_*, we find that

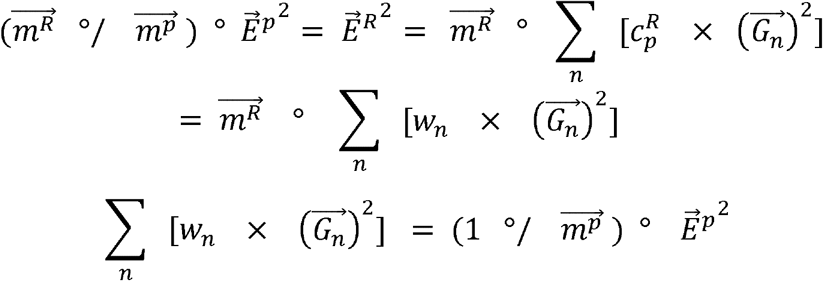

This equation can then be posed as a matrix equation

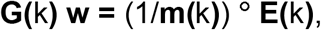

which can be solved using the non-linear least squares algorithm, as the GNBP weights should be non-negative. The non-linear least squares algorithm is run using the function *lsqnonneg* in MATLAB 2019a. In this paper, **G(**k) is a 5×3 matrix, **w** is a 3×1 vector, **m(**k) is a 5×1 vector, and **E(**k) is a 5×1 vector. The spectral calibration vector 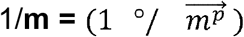 accounts for both the chromatic aberration 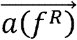 and the incident electric field 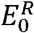.

In practice, we solve the de-mixing equation twice. We first solve the equation **G(**^1^**w) = (**^1^**E)**, with the spectral calibration set to unity, ^1^**m** = 1, as we don’t know the correct spectral calibration vector. For the 3 GNBP species, *n* = 1,2,3, we find pixels where the proportion, 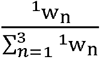, is between the 95^th^ and 100^th^ percentile for all pixels in the OCT volume (Figure S9a),. The average spectra of these pixels, 〈*G_n_*〉 is spectrally calibrated to yield pure particle spectra. The spectral calibration is then ^2^**m =** ^1^E °/ **<G_n_>**, which is used to recalibrate the electric field of the OCT volume to 1/(^2^**m)°/** (^2^**E)** (Figure S9b). We solve the de-mixing equation again **G(**^2^**w) =** 1/(^2^**m)** °/ (^2^**E)** to arrive at the final GNBP concentrations, ^2^**w**.

We apply the same spectral calibration ^2^**m** to all pixels in the OCT volume (Figure S9d), in spite of the fact that the incident electric field and chromatic aberration should be dependent on the depth for scattering media. In practice, this works because of the long depth-of-field of the lens, and the fact that the lymphatic vessels in mouse ear lie at very nearly the same depth in the mouse ear. This can be observed in Figure 4e,f,g. Within the depth-of-field, the GNBP spectra are indeed separable, even when not accounting for the change in spectra due to absorption and scattering (Figure S10). A depth-dependent chromatic aberration correction is also possible by localizing the focus in the OCT image^23^, but is not explored in this work.

### Algorithm Pipeline

The entire pipeline for converting an OCT interferogram to a de-mixed OCT image, with proportions of GNBPs, is shown in Figure S11. We first pre-process the interferogram by subtracting the DC apodization term. We then divide this result by the DC apodization term, factoring out the OCT light source spectral shape (citation). The modified interferogram is processed using a discrete-time short-time fourier transform (STFT) with gaussian window defined as:

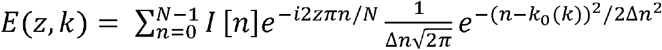

Where *n* is the wavenumber index of the spectrometer camera, *I*[*n*] is the interferogram, *N* = 1024 is the total number of spectrometer pixels, Δ*n* = 300 is the width of each spectral band, *k* is the index of the STFT spectral band, and *k*_0_ is the central wavenumber index of each spectral band. Each spectral band is shifted from the previous by a fixed interval allowing for overlap between spectral bands (Figure S11a), such that *k*_0_(*k*_*i*+1_) – *k*_0_(*k_i_*) = 181, allowing for an average overlap of 26.5 nm. Additionally, we apply standard OCT dispersion compensation techniques, by including a phase with square dependence on the wavenumber^24^. The STFT is masked using a speckle variance mask from the standard OCT image (Figure S11d), computed using a non-equispaced discrete discrete fourier transform^25^. The STFT spectra *E*(*z*, *k*) are corrected using the pre-measured falloff calibration curves (Figure S11g). The demixing algorithm is run on the spectrum of each pixel in the image, *E*(*k*) with the spectral calibration set to unity. The correct spectral calibration is computed (Figure S11j) and the algorithm is re-run with the new spectral calibration to yield the final result (Figure S11h).

### Chromatic Aberration Characterization

The chromatic aberration is measured using a capillary loaded with GNBP_1225_ at OD 5 (0.167 nM). The focus position is set to the very top of the capillary, which is observed when the common path image is visible at the top of the OCT image, from a strong reflection at the air-glass capillary interface. The focus is shifted to 4 positions above and 4 positions below the top of the glass capillary, in increments of 80 μm. The capillary is kept at the exact same position in the image after shifting the focus, but adjusting the reference arm length correspondingly. The data is spline interpolated at 10x of the original sampling density to yield smooth curves (Figure S8c). The chromatic aberration measurements are not utilized in the current pipeline, as they are implicitly compensated for by the spectral calibration.

### OCT Sensitivity Fall-off Calibration

The spectrally dependent sensitivity fall-off is measured using a capillary loaded with GNBP_1225_ at OD 5 (0.167 nM). The reference arm position is shifted, placing the capillary at 13 positions in the OCT image, that are 180 μm apart. The data is spline interpolated at 10x of the original sampling density to yield smooth curves (S7c), which are used to calibrate the STFT spectra from images.

### Principle Component Analysis

Principal Component Analysis (PCA) is used to analyze the STFT spectra in a lower dimensionality, and visualize the fact that the spectra for different GNBP species belong to distinct clusters that are separable in PCA space. PCA is a procedure for finding an orthogonal set of basis vectors for which the data is uncorrelated^15^. It is used here to reduce the 5-dimensional spectra to 3-dimensions for mapping to RGB images (Figure S1a) and to 2-dimensions for visualization in a scatter plot (Figure S1b). The singular values of the first three PCA components are 0.149, 0.026, and 0.005, thus justifying visualization using only the first two components.

The GNBPs are also separable even when the optical focus is placed at various positions relative to the capillary tube surface (Figure S12). The main effect the different focus positions is to induce a different chromatic aberration on the STFT spectra from pixels at the top of the capillary. Clusters representing various mixtures, coded by different colors (Figure S12b), occupy distinct positions in the 2D PCA plane.

We also show the PCA decomposition of the spectra, mapped to RGB for the 15 minute post-injection image (Figure S13). The color scheme is observed to resemble that of Figure 1a, as expected, showing that each GNBP channel consists of a distinct PCA decomposition.

### *In vitro* Imaging

Square glass capillary tubes (0.5 mm inner edge length and 0.1 mm wall thickness, VitroCom Inc., Mountain Lakes, NJ) were used to investigate the spectral de-mixing of GNBP_1225_, GNBP_1315_ and GNBP_1450_ *in vitro*. The OCT B-scan images of parallel capillary tubes containing pure and different molar mixtures of GNBP_1225_, GNBP_1315_ and GNBP_1450_ aqueous solutions, 1:0:0, 0:1:0, 0:0:1, 3:1:6, 1:3:6, 3:2:5, 2:3:5, 2:8:0: 8:0:2 and 0:8:2 from left to right in Figure 2a,b,c), with total concentration of OD 5 (0.167 nM). The focus in the de-mixed images in Figure 2a,b,c is 200 μm above the surface of the glass capillary. To analyze the spectral signal of each GNBP sample, a region of interest (ROI) of 420 μm × 71 μm was marked 5 pixels below the top of the first capillary tube (but located at the same height for all capillaries), in the acquired OCT B-scan images. We apply a 2×2 median filter to 2D images to reduce noise, bringing the effective lateral resolution to 12 μm, which is approximately equal to the optical resolution of the lens.

### Animal handling and *in vivo* imaging

All animal work was conducted in compliance with the guidelines of Stanford University’s Animal Studies Committee for the Care and Use of Research Animals and the experimental protocols were approved by the committee (APLAC protocol 27499). 6–8 weeks old female nude (nu^-^/nu^-^) mice (Charles River Inc., Wilmington, MA) were used for all *in vivo* experiments. During all imaging experiments, the mice were anesthetized by 2% isoflurane and placed on a 37 °C heating pad. The pinnae of the mice were attached to a mount by double-sided tape to minimize tissue motion.

To conduct the spectral de-mixing studies of OCT contrast agents GNBP_1225_, GNBP_1315_ and GNBP_1450_ *in vivo*, we subcutaneously injected 0.1 μL of OD 150 (5 nM) of PEGylated GNBP_1225_, GNBP_1315_ and GNBP_1450_ on the distal end of mouse pinnae at discrete locations which were least 1 mm apart from each other. The injections were conducted using a 2.5 μL microsyringe (Hamilton, Reno, NV) with a 31-gauge needle. The optical focus was placed 100 μm above the surface of the mouse ear, as lymph vessels tend to be 50-100 μm below the surface of the tissue. Three dimensional OCT images of the entire mouse ear, consisting of 6, 6×1 mm volumes, were acquired prior to injection, 15 min and 60 min after the 3 injections.

After performing de-mixing on the OCT volume, we apply a 2×2×2 median filter to 3D images to reduce noise, bringing the effective lateral resolution to 12 μm, which is approximately equal to the optical resolution of the lens.

We use the pre-injection image to remove the blood vessels in the post-injection image. The pre-injection was first registered to the post-injection images via monomodal 3d registration with translation and rotation degrees of freedom (Figure S14b). The registered pre-injection image was then spherically dilated and subtracted from the post-injection image (Figure S14c). This is successful in removing the majority of large blood vessels, leaving some edges which were incompletely removed, and some smaller blood vessels which are not removed. The remaining blood vessels are cleaned by hand in the xz plane of the tiff image (Figure S14d). Blood vessels can be distinguished from lymphatic vessels by extensive shadowing effects of the former, generating speckle variance signal that is observed in multiple layers throughout the mouse ear.

## Data availability

The data that support the plots within this paper are available in the main text and supplementary figures. Additional code and information are available from the corresponding authors upon request.

## Acknowledgments

This work was funded in part by grants from the United States Air Force (FA9550-15-1-0007), the National Institutes of Health (NIH DP50D012179), the National Science Foundation (NSF 1438340), the Damon Runyon Cancer Research Foundation (DFS# 06-13), Claire Giannini Fund, the Susan G. Komen Breast Cancer Foundation (SAB15-00003), the Mary Kay Foundation (017-14), the Skippy Frank Foundation, the Donald E. and Delia B. Baxter Foundation, a seed grant from the Center for Cancer Nanotechnology Excellence and Translation (CCNE-T; NIH-NCI U54CA151459), and a Stanford Bio-X Interdisciplinary Initiative Seed Grant (IIP6-43). A.D.Z. is a Chan Zuckerberg Biohub investigator and a Pew-Stewart Scholar for Cancer Research supported by The Pew Charitable Trusts and The Alexander and Margaret Stewart Trust. A.D.Z is a Chan Zuckerberg Biohub investigator and a Pew-Stewart Scholar for Cancer Research supported by The Pew Charitable Trusts and The Alexander and Margaret Stewart Trust. We would like to acknowledge Yonatan Winetraub for discussion regarding analysis of the results in this work.

## Author Contributions

E.Y., P.S., A.D.Z. conceived of the idea of multiplexed OCT nanoparticles. E.Y. developed the methodology, spectral processing technique, calibration, and code. P.S. created the gold nanobipyramids. E.Y., P.S., and S.S. acquired the data. E.Y. analyzed the data and created the figures. E.Y., P.S., A.D.Z. aided with interpretation of the results. E.Y., P.S., A.D.Z wrote the manuscript. E.Y., P.S., A.D.Z. aided in editing the manuscript and figures.

