## Supplementary Information for "A Spectral De-mixing Model for Triplex *In Vivo* Imaging of Optical Coherence Tomography Contrast Agents"

for

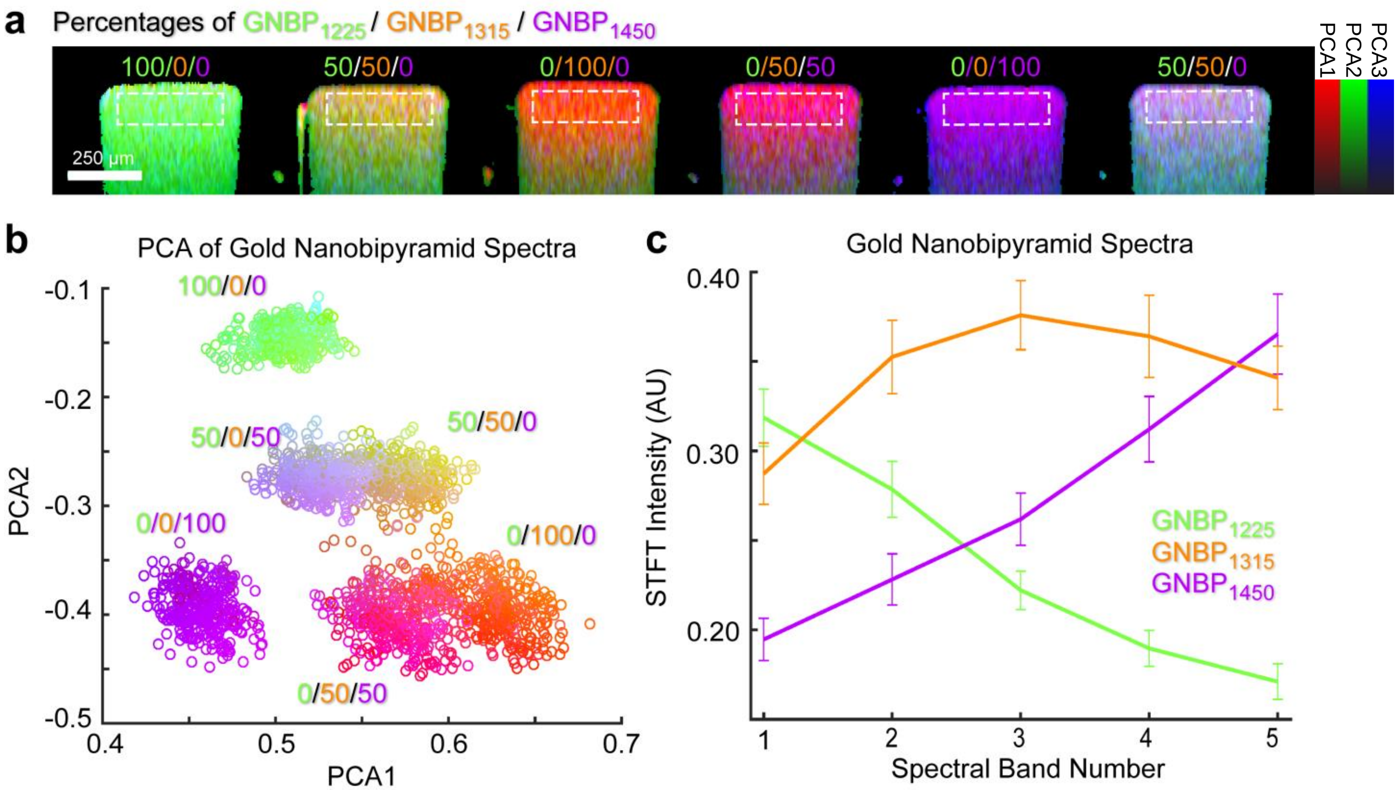


**Figure S1 PCA Analyis of GNBP STFT spectra. a** The original 5-dimensional STFT spectra of various mixtures of GNBP (all at OD 5 or 0.167 nM) are reduced to 3 PCA dimensions and visualized as RGB colors, with red: PCA1, green: PCA2, and blue: PCA3. From left to right, capillaries contain pure $GNBP_{1225}$, 50/50% mixture of $GNBP_{1225}$ and $GNBP_{1315}$, pure $GNBP_{1315}$, 50/50% mixture of $GNBP_{1315}$ and $GNBP_{1450}$, pure $GNBP_{1450}$, and 50/50% mixture of $GNBP_{1225}$ and $GNBP_{1450}$. Scale bar: 250 µm. **b** Scatter plot of 2 leading PCA components, taken from the ROI’s indicated by dotted white boxes in **a**. Each GNBP mixture occupies a distinguishable cluster in PCA space. **c** Five-dimensional STFT spectra of pure particles, $GNBP_{1225}$, $GNBP_{1315}$, and $GNBP_{1450,}$ from ROIs indicated by the white dotted box in **a**.

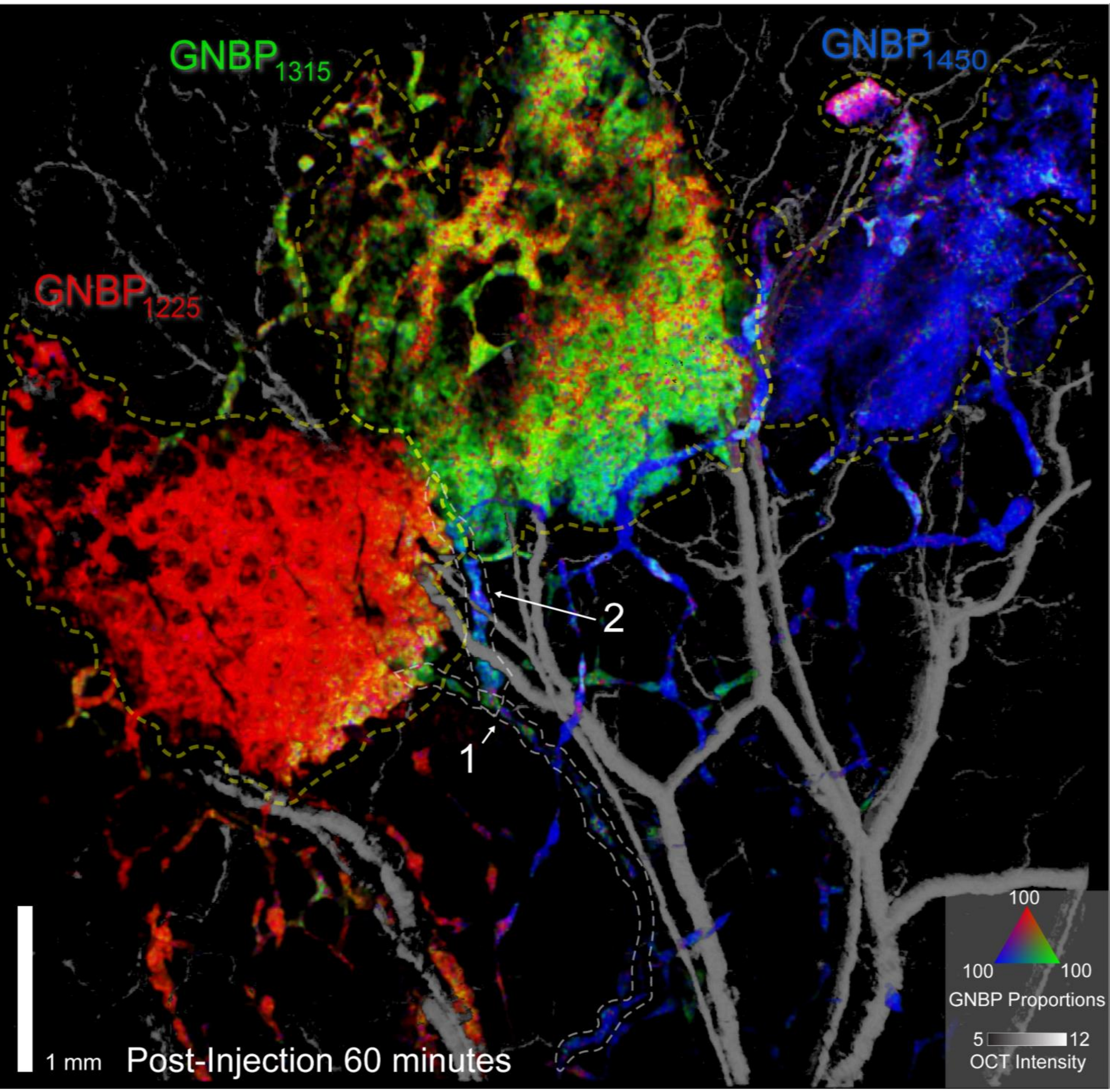


**Figure S2 De-mixed image 1 hour post-injection.** De-mixed image of OCT volume of mouse ear 1 hour after injection of $GNBP_{1225}$, $GNBP_{1315}$, and $GNBP_{1450}$. OCT volume is gated by speckle-variance to only show flow – blood vessels and lymph vessels. Compared to the de-mixed image 15 minutes after injection, injections sites are observed to have widened in area, due to subcutaneous diffusion of GNBPs. Injection sites are labeled by dotted yellow lines, and two lymphatic vessels of interest, lymph 1 and lymph 2, are labeled by dotted white lines

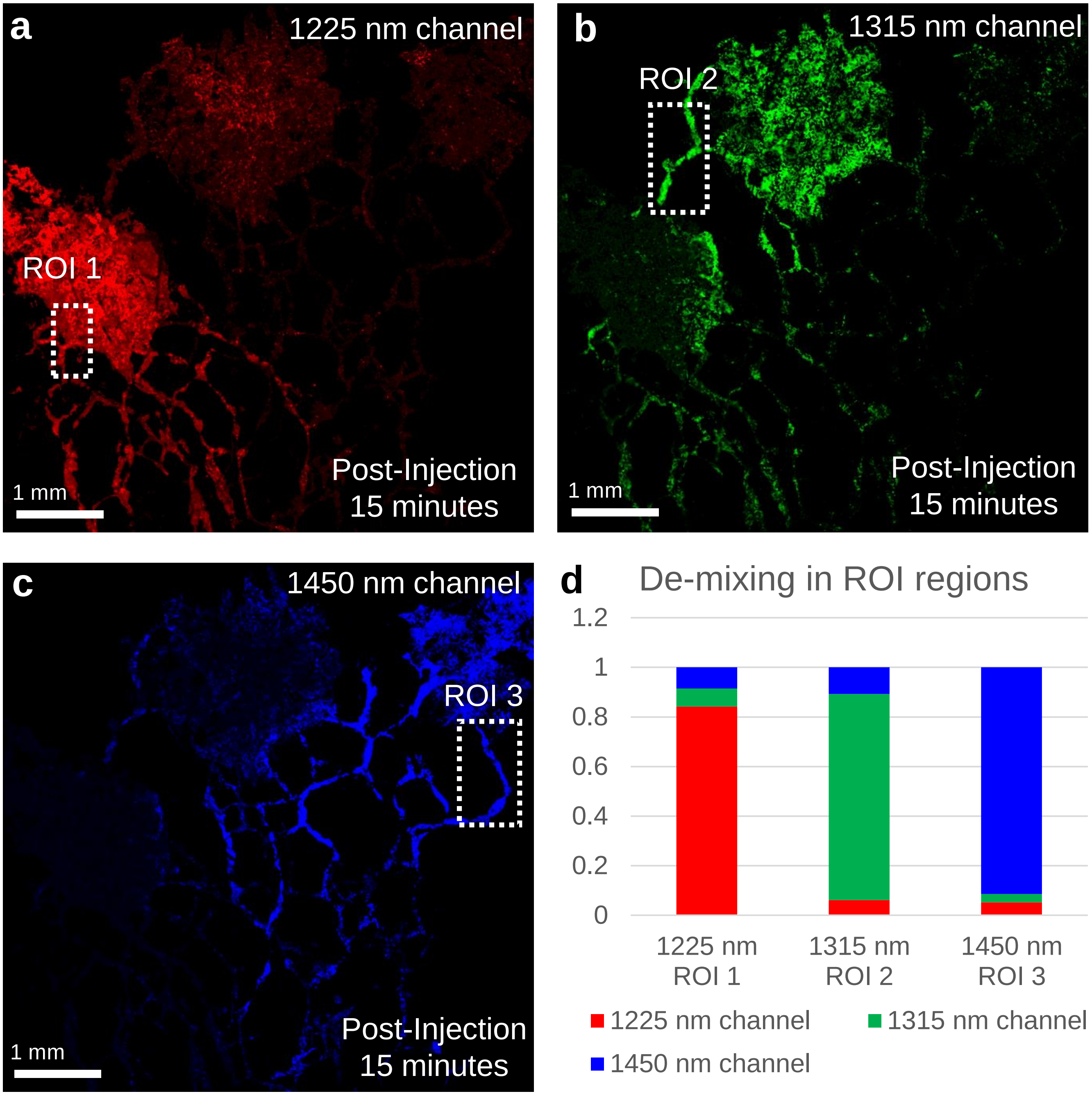


**Figure S3 Individual channels for de-mixed 15 minutes post-injection image. a** 1225 nm channel. **b** 1315 nm channel. **c** 1450 nm channel. **d** Assessing the accuracy of the de-mixing by measuring the predicted GNBP proportions in selected regions of high single GNBP purity. In each channel, a rectangular ROI is selected, and the average GNBP proportion is reported. Numerical results are shown in **Table S1.**

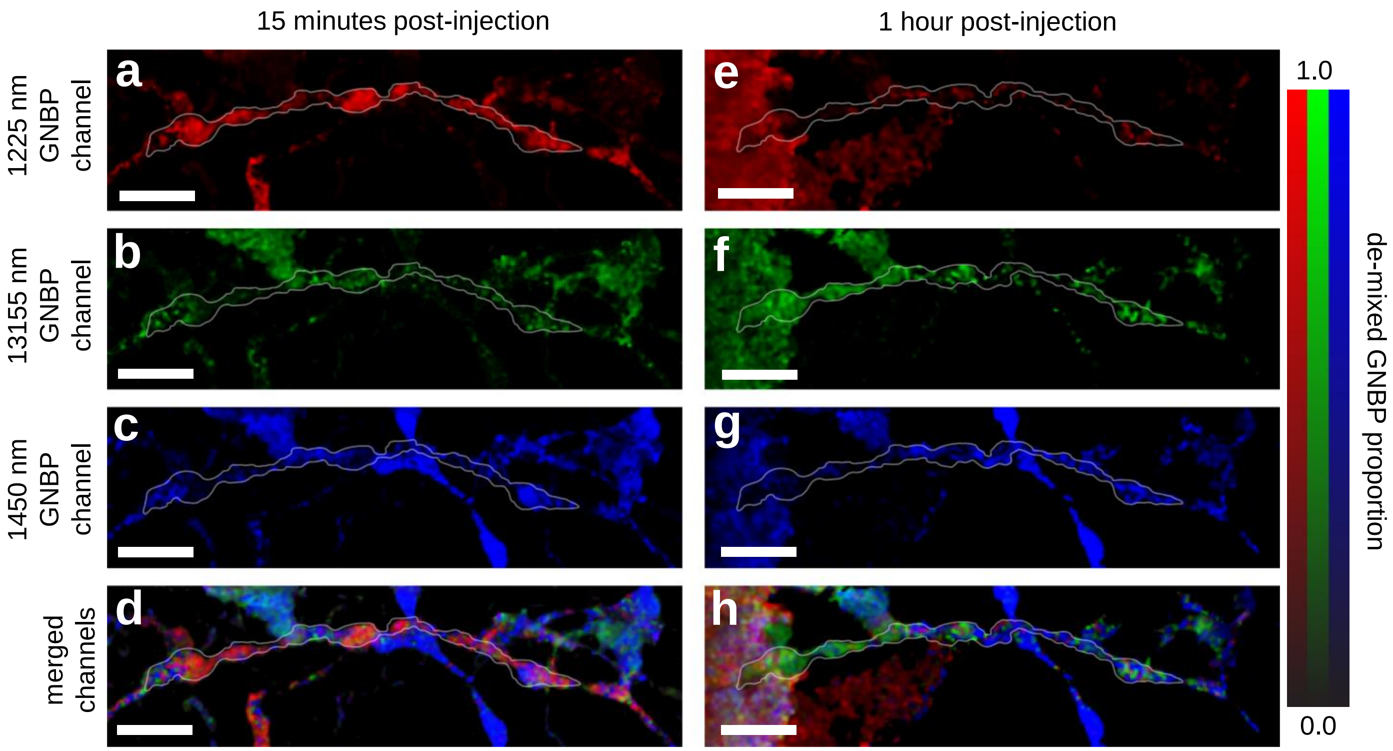


**Figure S4 Comparison of lymph 1, 15 minutes and 1 hour post-injection.** De-mixed images of lymph 1 (**Figure 3**), after injection of $GNBP_{1225}$, $GNBP_{1315}$, and $GNBP_{1450}$. Images from each channel are shown for 15 minutes post-injection in **a** 1225 nm GNBP, **b** 1315 nm GNBP, **c** 1450 nm GNBP, and **d** merged channels. Similarly, for 1 hour post-injection, image channels are shown for **e** 1225 nm GNBP, **f** 1315 nm GNBP, **g** 1450 nm GNBP, and **h** merged channels. We observe that the GNBP composition of lymph 1 changes significantly by the 1 hour post-injection timepoint. There is a notable decrease in the proportion of $GNBP_{1225}$, and an increase in $GNBP_{1315}$ throughout the vessel. White outline around lymphatic vessel boundaries shows the outline of the lymph1. Scale bars: 400 μm.

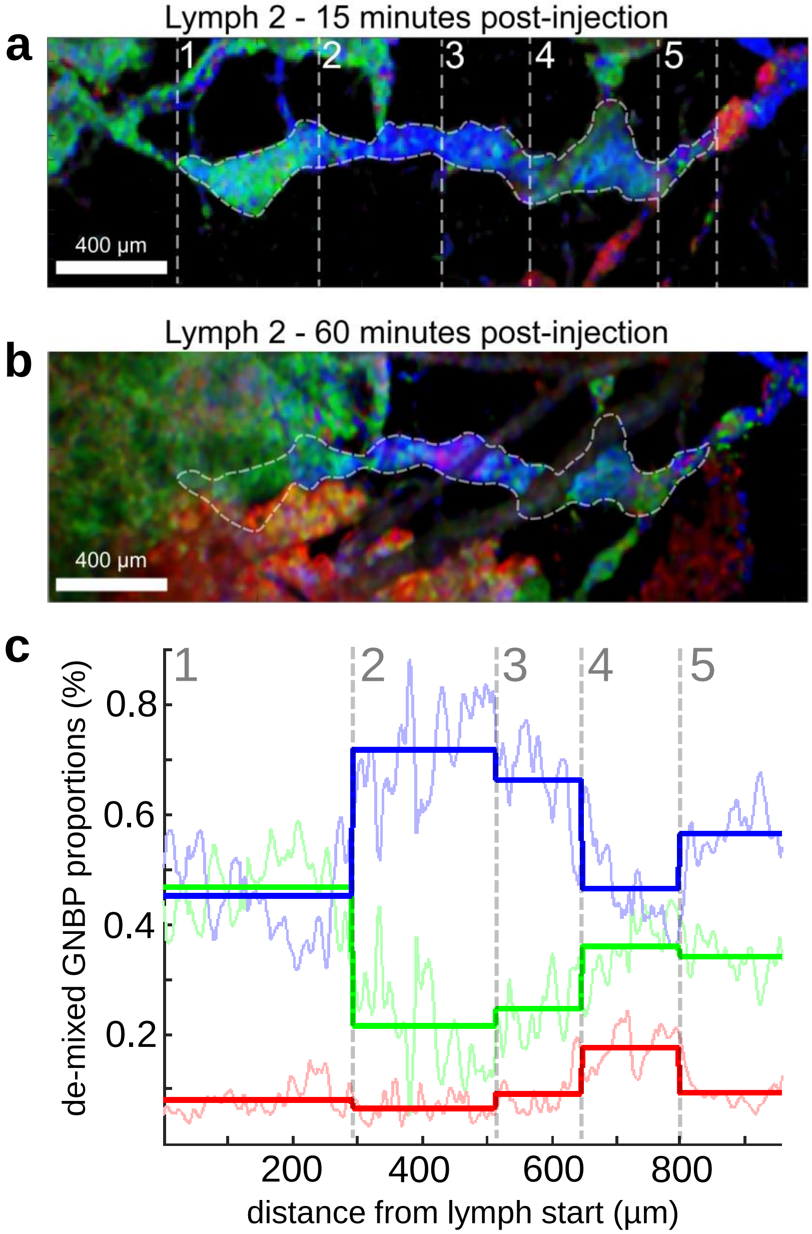


**Figure S5 De-mixed image of lymph 2. a** Lymph 2, 15 minutes after injection. Segmented lymph vessel is outlined with a white boundary and divided into 5 sections. **b** Lymph 2, at 60 minutes after injection. The GNBP composition of lymph 2 does not change significantly after 45 minutes, indicating that in spite of continuous pulsed lymph flow reported in literature, existing GNBP proportions are not changing significantly. **c** De-mixing of Lymph 2, 15 minutes after injection as a function of distance from the start of the lymph vessels, along a path following the lymphatic vessel curvature. Solid lines indicate average GNBP proportion of section of the lymph. Dimmed color lines show the data of the raw GNBP proportions. Scale bars: 400 µm

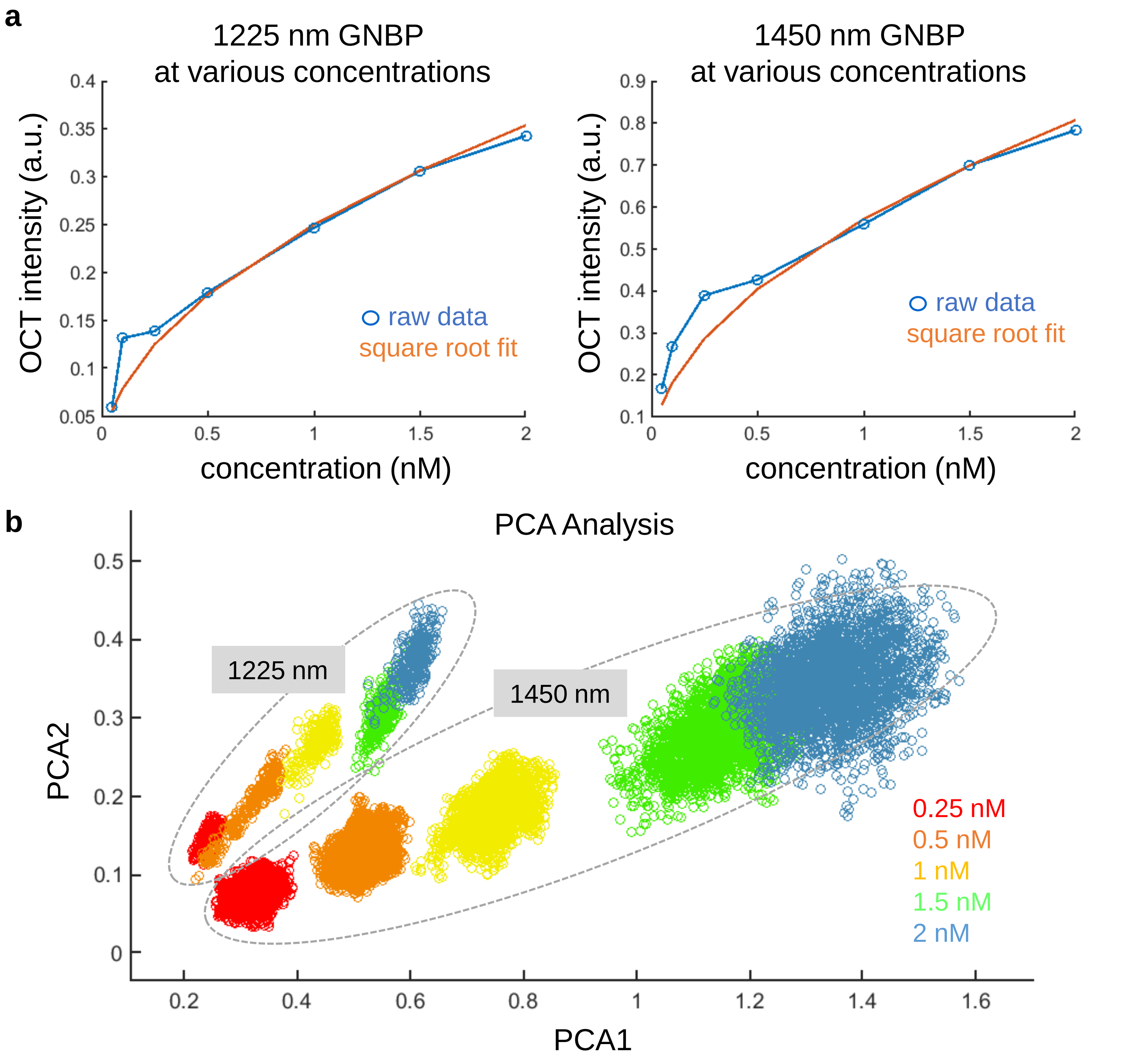


**Figure S6 Analysis of $GNBP_{1225}$ and $GNBP_{1450}$ at various concentrations.** **a** Measurements of $GNBP_{1225}$ and $GNBP_{1450}$ at concentrations of 0.05, 0.1, 0.25, 0.5, 1, 1.5, and 2 nM. As expected, at larger concentrations, the OCT intensity increases as the square root of the concentration, following the average of the Rayleigh distribution. **b** Principal Component Analysis (PCA) of the spectra for various concentrations. The two PCA components PCA1 and PCA2 are the leading principal components, and capture the shape of the spectra of the GNBPs. Plotting PCA1 and PCA2 on a scatter plot shows that increasing concentration yields a linear trend in PCA space, indicating that spectra shape does not change significantly as the concentration is increased.

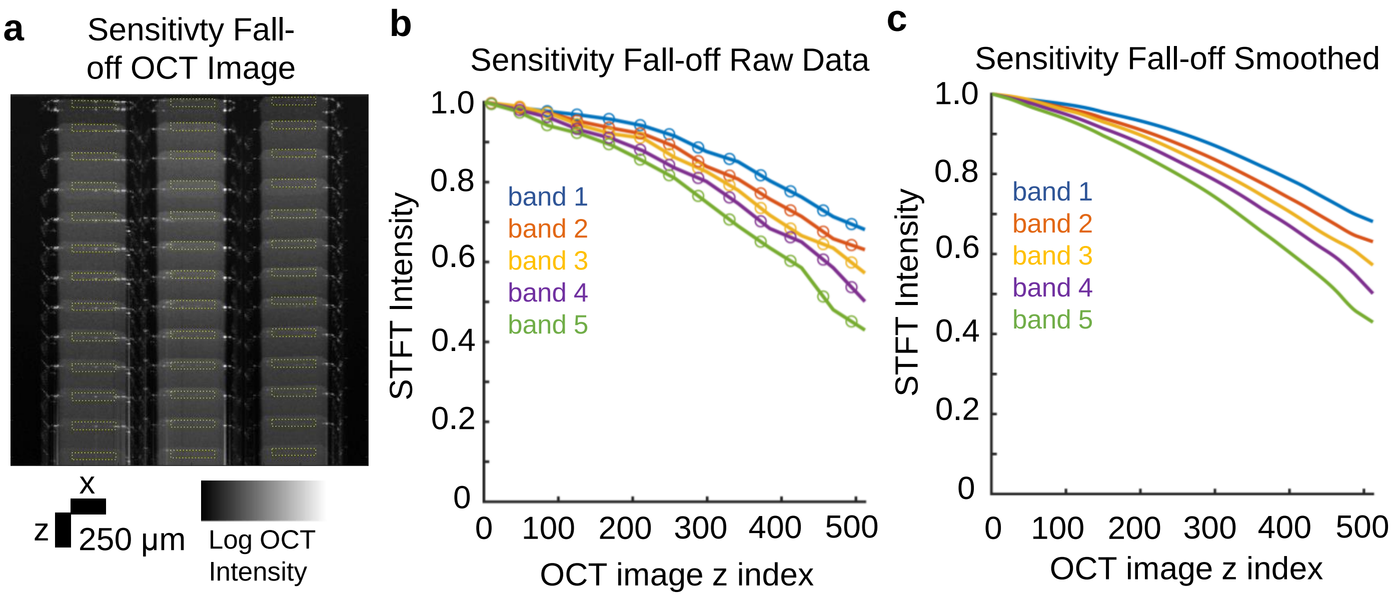


**Figure S7 OCT Sensitivity Fall-off measured for Thorlabs Telesto II system.** **a** Various OCT images of capillaries loaded with $GNBP_{1225}$ (OD 5 or 0.167 nM), as the reference arm position is shifted, while keeping the focus at a fixed position in the image. **b** Data points (open circles) from each of the reference arm measurements in **a,** where the average STFT intensity is taken from an ROI at the top of each capillary (dotted yellow rectangles). The interporlated curves are also shown as solid colored lines for each band. **c** 'Lowess' smoothed curves (span 100) of interpolated data from **b**. This interpolated data is used as the reference fall-off calibration curves in the pipeline **S2**. Scale bars are 250 µm in both the x and z directions.

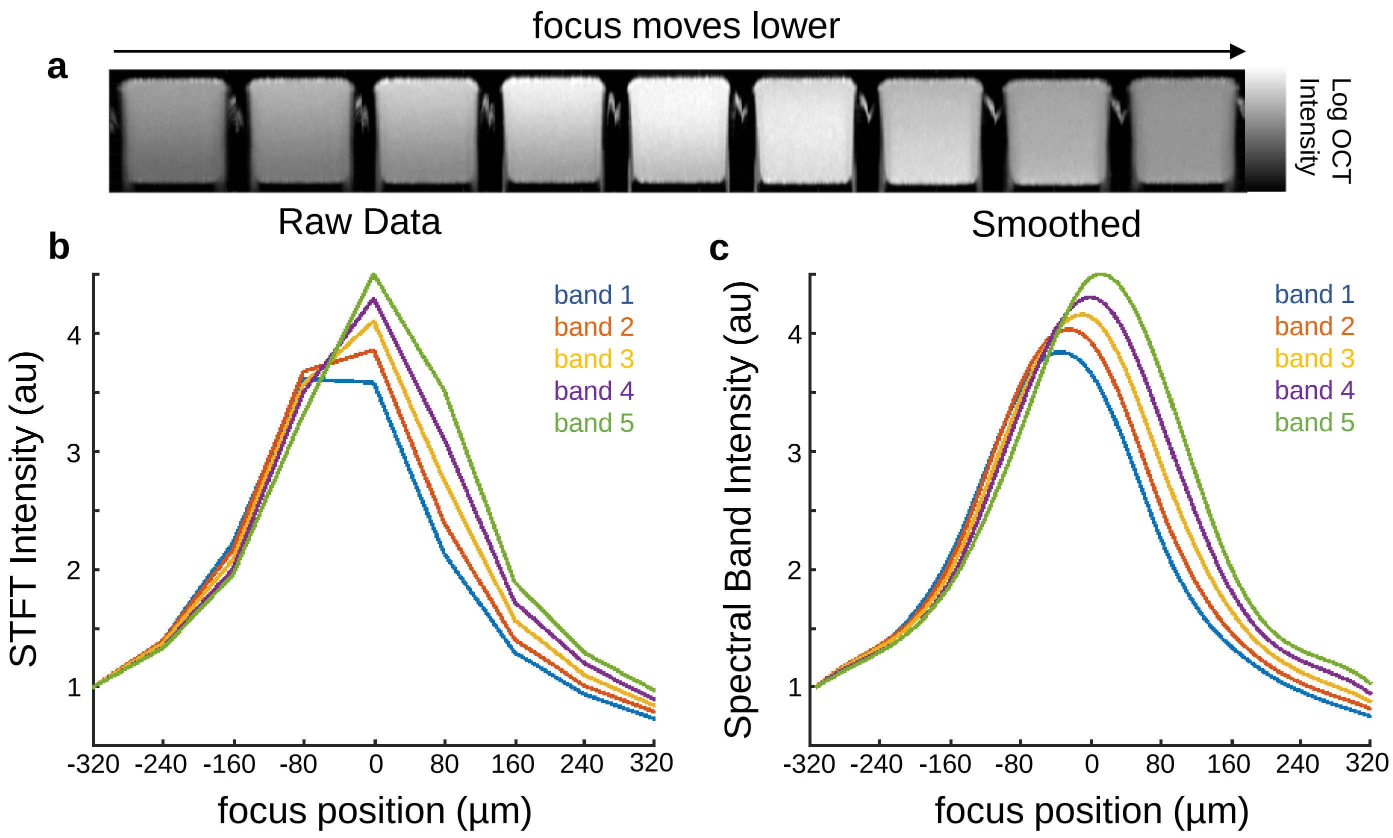


**Figure S8 Chromatic Aberration of LSM03 lens for Thorlabs Telesto II system.** **a** OCT images of capillaries loaded with $GNBP_{1225}$ (OD 5 or 0.167 nM), as the focus position moves lower in the image in increments of 80 µm. **b** OCT intensity of the 5 spectral bands (various colors), as the focus position moves lower in the image**.** Each spectral band peaks at a different location, showing the chromatic aberrations characteristic of the particular lens used in the experiment. **c** Smoothed spectral bands from **b,** showing more clearly, the different peaks of each spectral band.

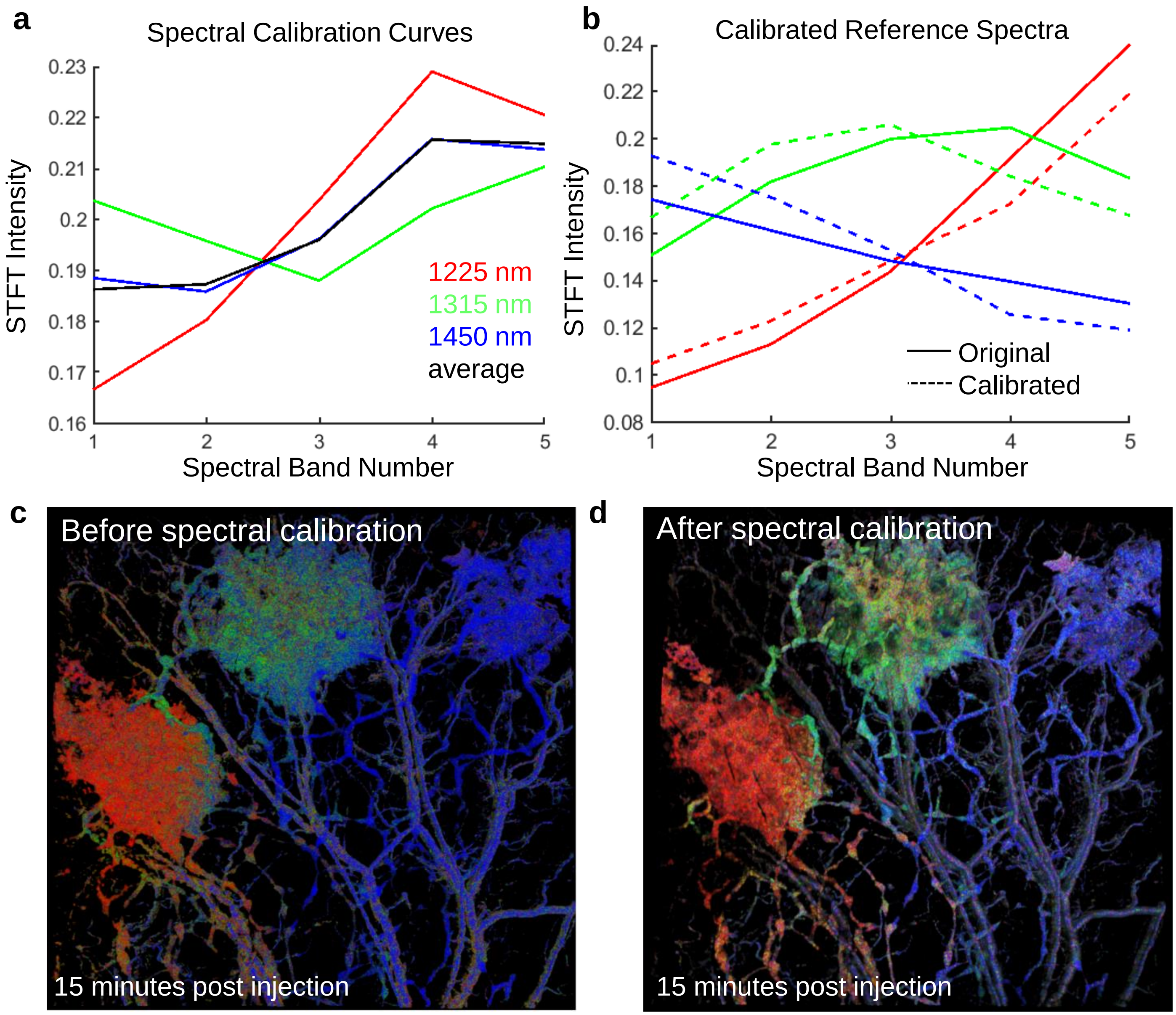


**Figure S9 Calculation of Spectral Calibration Curves and Before/After.** **a** Spectral calibration curves calculated from the 95 percentile and and above of each channel. The average spectral calibration curve which is used in for calibration is shown in black**.** **b** The calibrated spectra are shown as dotted lines, after using the calibration in **a.** The original spectra are shown as solid lines. **c** De-mixed **i**mage of in-vivo injection of 3 GNBP before spectral calibration. The spectra of the entire image is red shifted (towards longer wavelengths), particularly noticeable in the blood vessels, which are colored blue (the color $GNBP_{1450}$). **d** De-mixed image after spectral calibration. Blood vessels have a more neutral color, indicating, on average, equal proportions of the 1225, 1315, and 1450 nm GNBP channels.

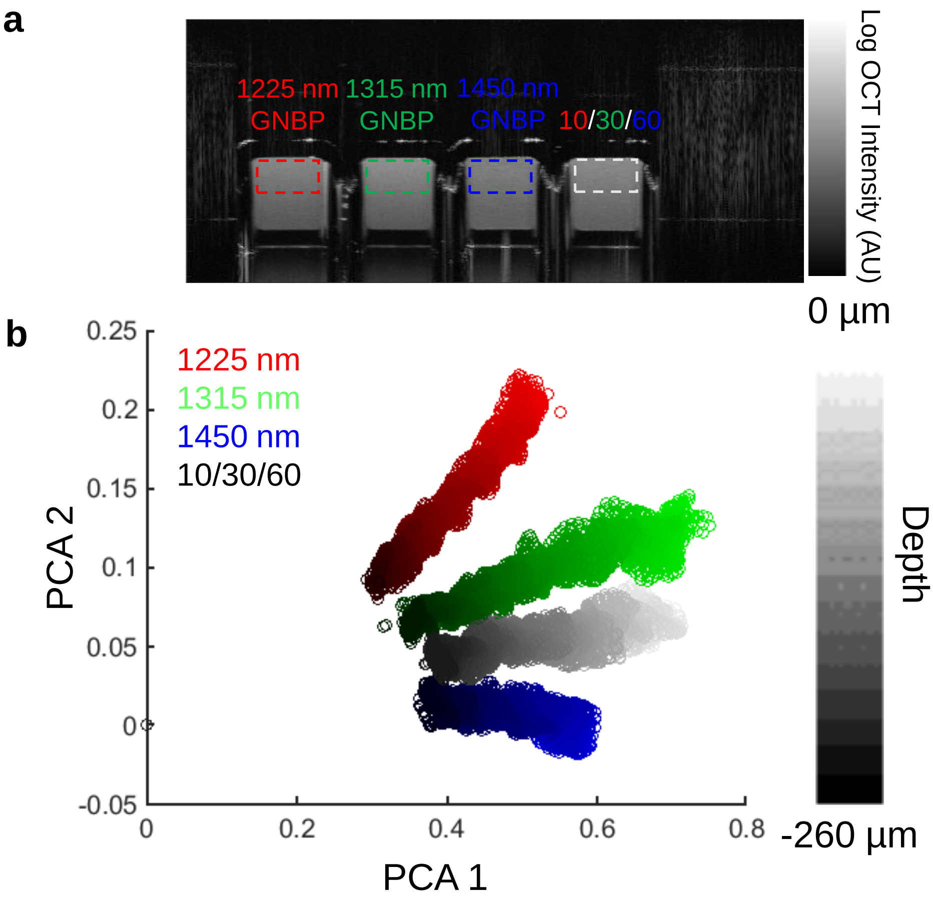


**Figure S10 Analysis of GNBP spectra as a function of depth.** **a** OCT image of $GNBP_{1225}$, $GNBP_{1315}$, and $GNBP_{1450}$ at OD 5, or 0.167 nM, as well as a mixture of 10%/30%/60%. **b** STFT spectra are measured from the top of capillaries in **a** (dotted ROI's), and Principal Component Analysis (PCA) is performed. The two PCA components PCA1 and PCA2 are the leading principal components, and capture the shape of the spectra of the GNBPs. Various data points are plotted from various depths. The depth is encoded by the intensity of the pixels. The results show that spectra are separable in PCA space, and allow for quantitative de-mixing. Additionally, the points from each capillary show a linear trend in PCA space, showing that the spectrum shape is maintained as a function of depth.

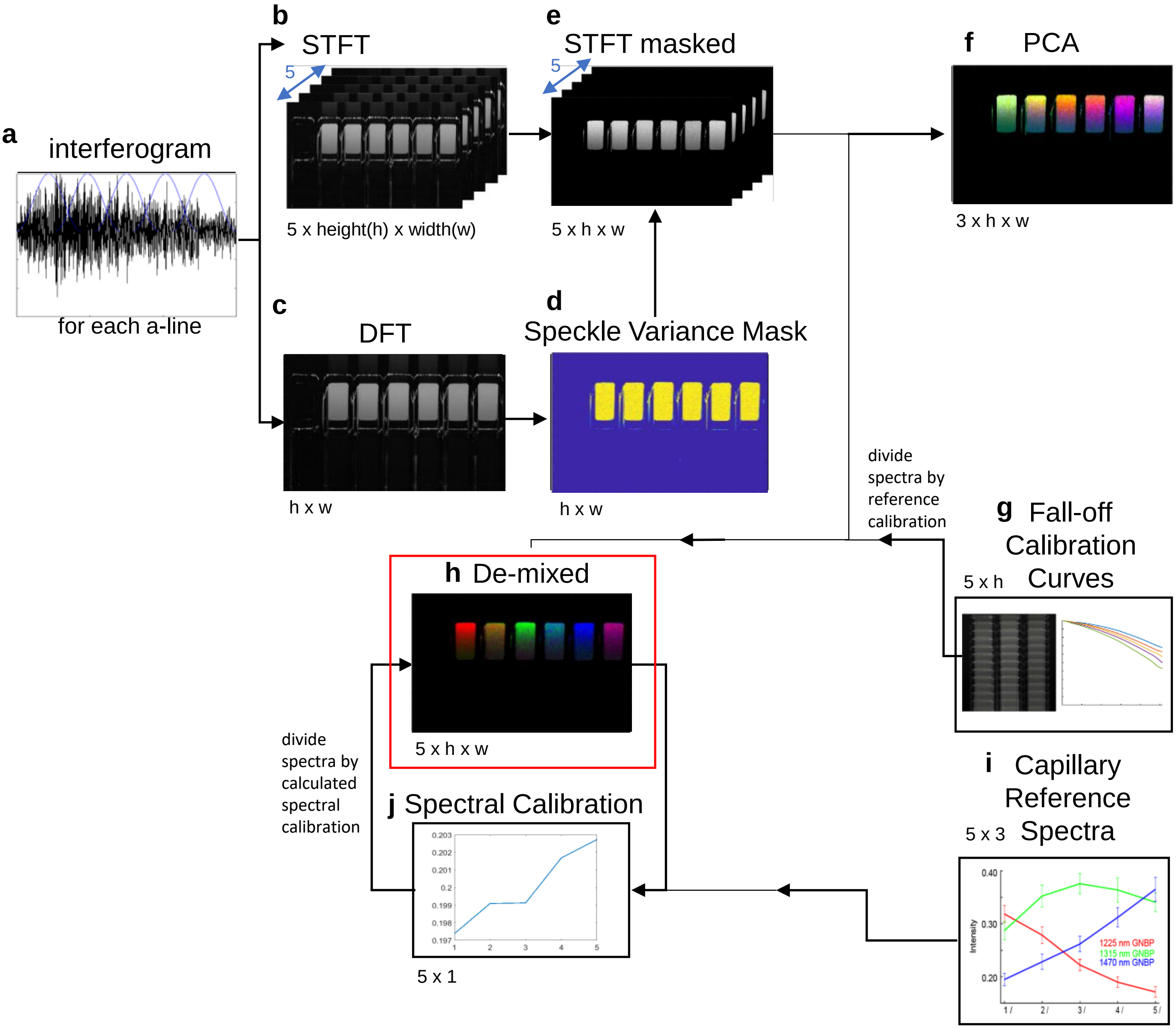


**Figure S11 Pipeline for converting OCT interferogram to a demixed OCT image with proportions of GNBPs.** **a** Single a-line interferogram obtained from camera. Five spectral bands are shown in blue. **b** Short-time fourier transform of interferogram in **a**, yielding 5 images, **c** Discrete fourier transform of interferogram in **a** is also taken. **d** Speckle-variance mask is calculated from the discrete fourier transform in **c.** **e** STFT is masked using speckle variance mask from **d.** **f** Principal component analysis can be performed on **e** to visualize and analyze the spectra produced by the STFT. **g** OCT sensitivity fall-off curves are measured and smoothed and used to calibrate the masked STFT image from **e.** **h** A demixed OCT image is produced using optimization of non-zero least squares on a per pixel basis. **i** Reference spectra are the experimentally measured reference spectra of $GNBP_{1225}$, $GNBP_{1315}$, and $GNBP_{1450}$. **j** The reference spectra are used to produce a spectral calibration, which is used to eliminate chromatic aberration, and other spectrally aberrant effects such as attenuation. **h** The de-mixed spectra are calibrated using the spectra calibration and then non-zero least squares optimization is performed again. Steps **h,I,j** are performed in two iterations to first determine the spectral calibration, and then to implement it.

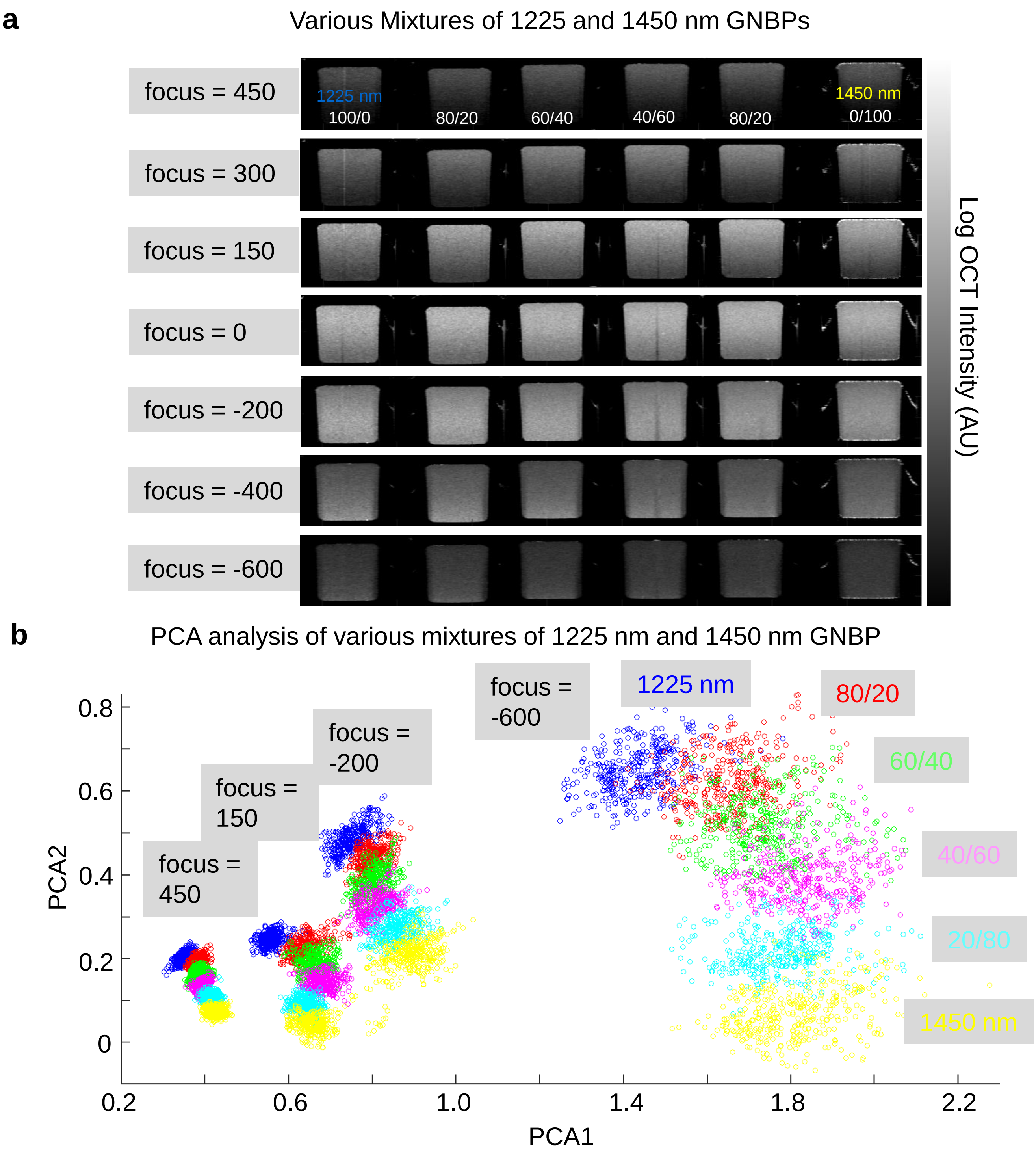


**Figure S12 Analysis of various mixtures of GNBP$_{1225}$ and GNBP$_{1450}$.** **a** Capillaries containg various mixtures of GNBP$_{1225}$ and GNBP$_{1450}$ (OD 5 or 0.167 nM). From left to right, the percentages of 1225 / 1450 nm GNBPs is 100/0, 80/20, 60/40, 40/60, 0/100. Additionally, each dataset is taken at various focus positions. **b** Principal Component Analysis (PCA) of the spectrum from the top of each capillary. The two PCA components PCA1 and PCA2 are the leading principal components, and capture the shape of the spectra of the GNBPs. Plotting PCA1 and PCA2 on a scatter plot shows strong separability each spectral cluster, both for various mixtures, and at various focal depths.

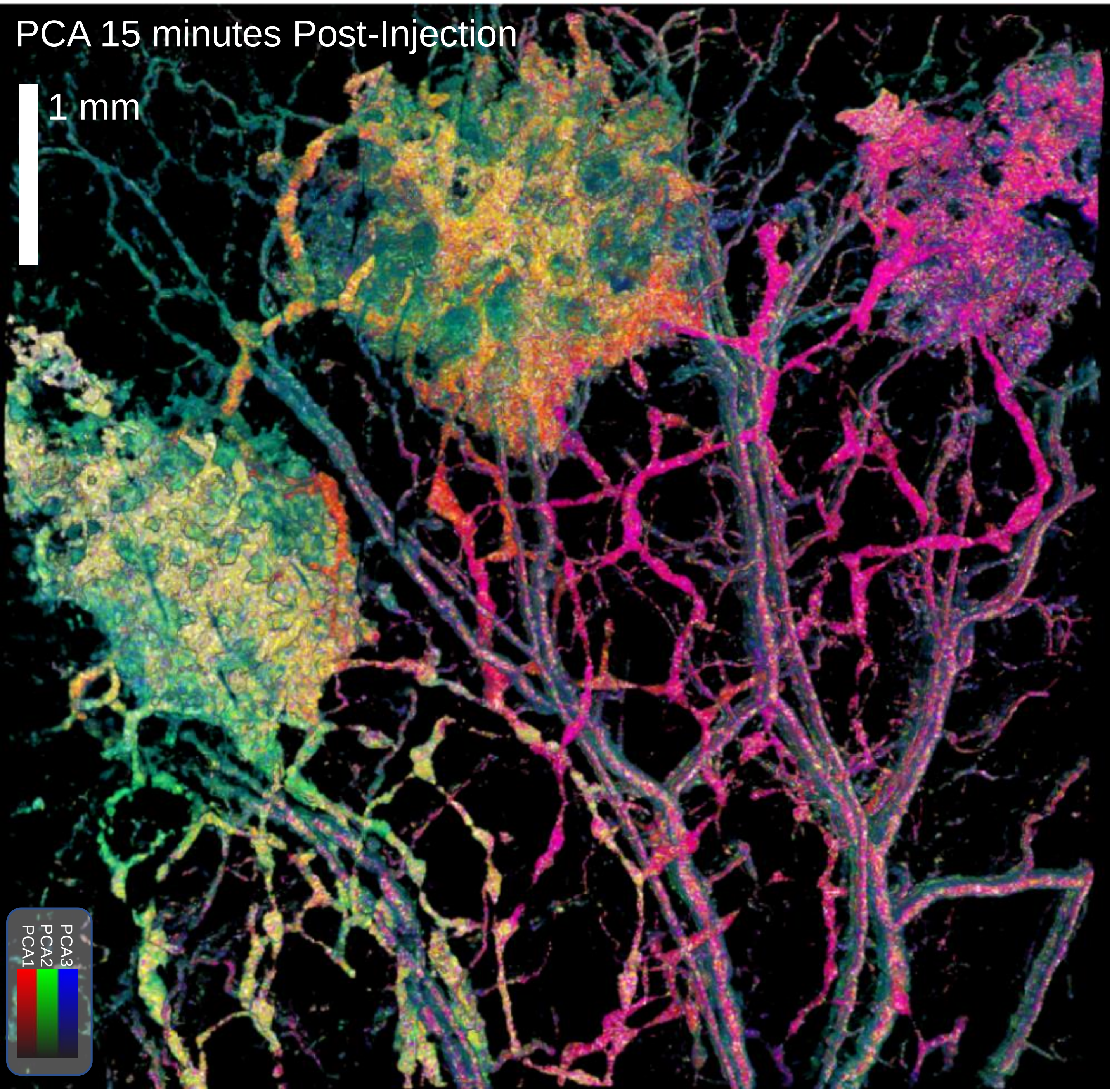


**Figure S13 3 component PCA of 15 minute post-injection image.** Principal component analysis (PCA) of spectra from OCT volume of mouse ear 15 minutes after injection of $GNBP_{1225}$, $GNBP_{1315}$, and $GNBP_{1450}$. OCT volume is gated by speckle-variance to only show flow – blood vessels and lymph vessels. Short-time fourier transform is applied to the raw interferometric data to produce a 5 band spectral image, which is reduced 3 components via PCA, and then mapped to colors RGB.

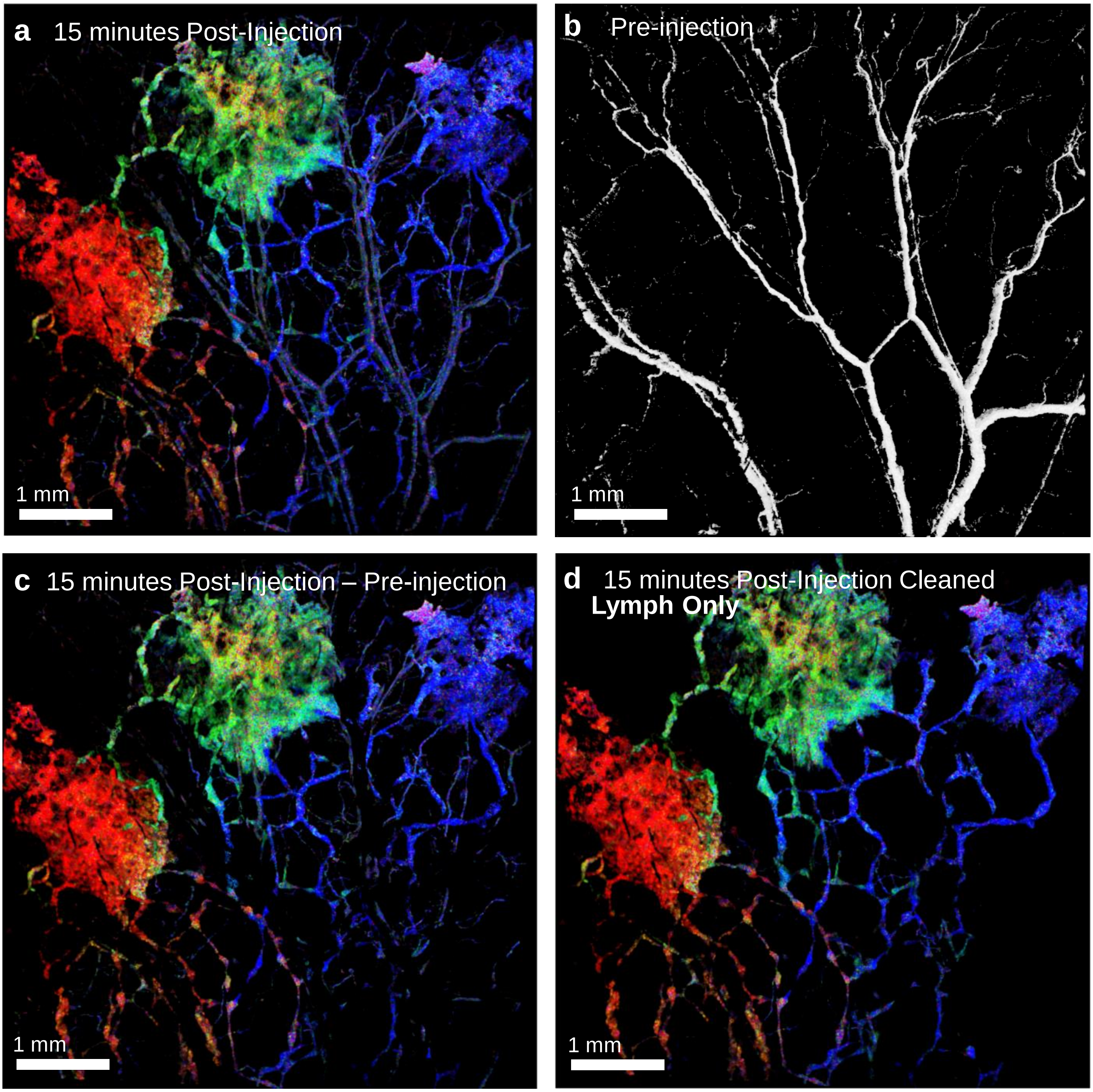


**Figure S14 Cleaning of images of blood vessels.** **a** De-mixed image of OCT volume of mouse ear 15 minutes after injection of $GNBP_{1225}$, $GNBP_{1315}$, and $GNBP_{1450}$. OCT volume is gated by speckle-variance to only show flow – of blood vessels and lymph vessels. **b** Pre-injection image gated by speckle-variance, only showing blood vessels, as GNBP have not yet entered lymph vessels. **c** De-mixed 15 minute post-injection image with blood vessels removed by subtraction with blood vessels from **b.** Prior to subtraction, blood vessels from **b** are spherically dilated by 5 pixels and registered to **a.** **d** De-mixed 15 minutes post-injection image after manually clearning and subtraction of pre-injection blood vessels. Manually cleaning is necessary as subtraction of blood vessels in **c** removes majority of large blood vessels, but leaves uncleaned edges as well as many smaller blood vessels. Blood vessels can be identified as they attenuate light more significantly compared to lymph vessels, and create shadowing throughout the entire depth of the mouse ear.

| | 1225 nm channel proportion | 1315 nm channel proportion | 1450 nm channel proportion | Average de-mixing error |
|---|---|---|---|---|
| $GNBP_{1225}$ ROI 1 | 0.8417±0.082 | 0.0734±0.044 | 0.0849±0.056 | 0.1053±0.061 |
| $GNBP_{1315}$ ROI 2 | 0.0614±0.033 | 0.8316±0.048 | 0.107±0.074 | 0.1123±0.052 |
| $GNBP_{1450}$ ROI 3 | 0.0523±0.026 | 0.0341±0.021 | 0.9136±0.063 | 0.0576±0.036 |

**Table S1 Numerical results from de-mixing of ROIs of high GNBP purity from 15 minutes post-injection.** Average demixed GNBP proportions from ROIs 1,2,3 in **S3.** Errors represent one standard deviation from the mean. This shows that the average error is 0.1053 for $GNBP_{1225}$, 0.1123 for $GNBP_{1315}$, and 0.0576 for $GNBP_{1450}$, higher than the average error of about 0.07 (**Fig 2e**) for de-mixing in capillaries, in-vitro. An explanation for the reduced error for $GNBP_{1450}$ compared to the other GNBPs is that the raw spectra were naturally red-shifted (**S3c**), and the spectral calibration was insufficient to fully balance the spectra, leading to a higher predicted proportion of $GNBP_{1450}$.
